# The Paipu framework enables creation of a large-scale mammalian cancer transcriptomics atlas

**DOI:** 10.64898/2026.05.14.725161

**Authors:** Bria S. Smith, Leslie A. Smith, Geesa Daluwatumulle, Nathan Glen, Ji-Hyun Lee, James A. Cahill, Kiley Graim

**Affiliations:** Department of Computer and Information Science and Engineering, University of Florida, 1889 Museum Rd, Gainesville, 32611, Florida, USA; Department of Biostatistics, University of Florida, 2004 Mowry Rd, Gainesville, 32603, Florida, USA; Department of Environmental Engineering Sciences, University of Florida, 365 Weil Hall, Gainesville, 32611, Florida, USA; UF Genetics Institute, University of Florida, 2033 Mowry Rd, Gainesville, 32610, Florida, USA; UF Health Cancer Center, University of Florida, 2033 Mowry Rd, Gainesville, 32610, Florida, USA

**Keywords:** comparative genomics, pan-cancer, pipeline, atlas

## Abstract

A plethora of studies have identified shared molecular mechanisms involved in tumor development across humans and other mammalian species. While these two-species analyses advance understanding of human disease, extending analysis across many species would provide evolutionary insight into molecular mechanisms driving human cancers. However, this expansion requires knowledge transfer and harmonization across species. Genomic differences between species, including variation in genome annotation quality, have historically hindered multi-species large-scale atlas creation. To overcome these challenges, we present Paipu, a comprehensive pipeline designed to streamline querying, preprocessing, harmonization, and retrieval of large-scale RNA-seq data and associated metadata from the NCBI Sequence Read Archive (SRA). Paipu facilitates multi-species analysis by creating a harmonized atlas from user-defined search terms and species. It consists of three components: reference genome preparation, SRA metadata retrieval, and RNA-seq data processing. We apply Paipu to 188 cancer-related terms in 240 non-human mammalian species, creating a harmonized atlas of 51,561 RNA-seq samples spanning 18 species and 83 cancers. This pan-mammalian pan-cancer atlas enables myriad comparative genomics analyses that leverage genetic variation to better understand rare human cancers. As such, Paipu serves as a resource for cross-species cancer genomics and supports atlas creation for any set of species and search terms.

**Graphical Abstract:** 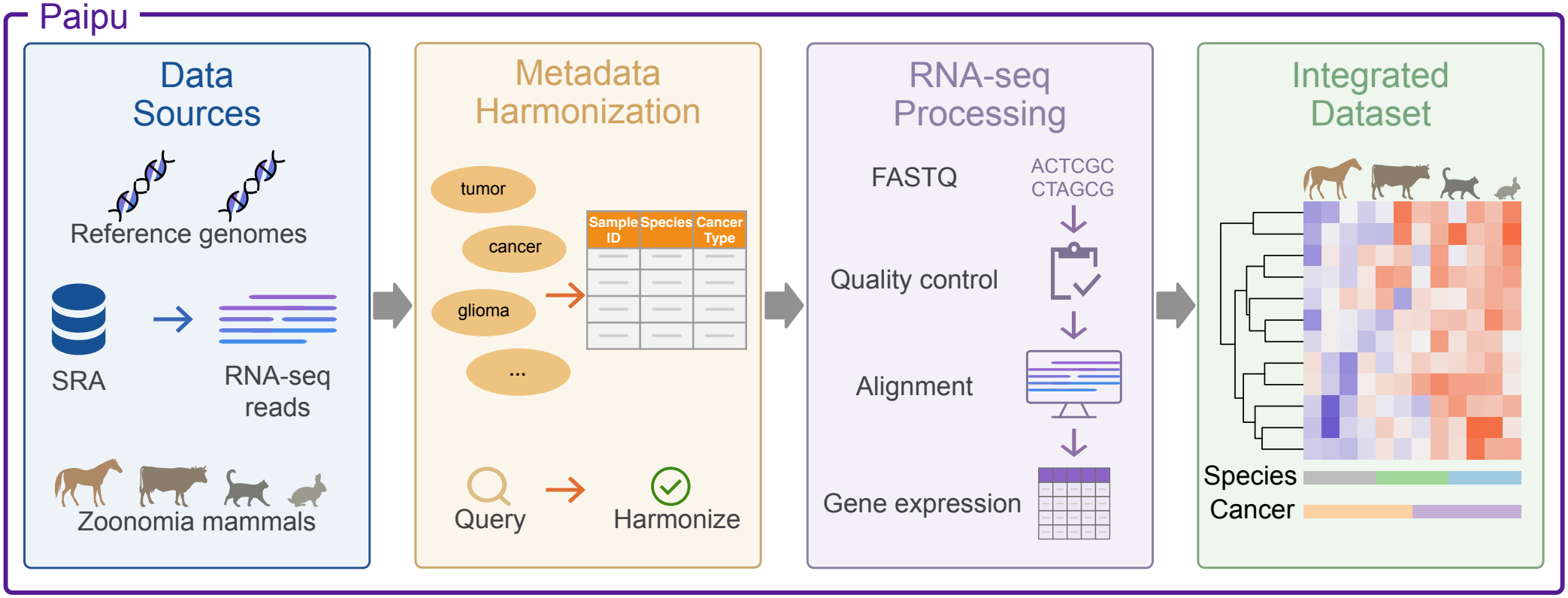

## Introduction

Advancements in high-throughput sequencing technologies have generated vast amounts of genomic data. As of Jan 01, 2026, *>*49 million BioSamples have been uploaded to the Sequence Read Archive. More than 8 million of these samples (~16%) were uploaded in 2025 alone. This large-scale data provides the opportunity for cross-species analyses on an unprecedented scale. Unfortunately, differences in reference genomes, gene annotations, and sequencing technologies have posed significant barriers to large-scale many-species data harmonization efforts. While databases such as the Sequence Read Archive (SRA) [45] store over 25 petabases of genomic data across more than 14 million sequencing runs [35], processing these data in a manner that allows for cross-species analysis is computationally intensive and requires comparative genomics expertise.

Previous comparative cancer genomics studies involving humans plus one other species have revealed key insights into disease mechanisms and evolutionary relationships. For example, studies have investigated cancer resistance in elephants [4], many spontaneous cancers in dogs [11, 26, 27, 43, 42], fibropapillomatosis in sea turtles [20], and genomic adaptations associated with cancer resistance in naked mole rats [61]. In addition, phylogenetic analyses identified 40 human cancer genes as being under positive selection in mammals [65]. While these studies have accelerated scientific understanding of human cancers, scaling these approaches to larger-scale studies presents significant technical challenges. Low-quality and poorly annotated genome assemblies can result in misalignment and inaccurate ortholog mapping [37, 40, 29, 55]. These and other challenges limit the potential of many-species analyses.

Despite these limitations, the benefits gained from increasing the number of species used in analyses are substantial. Integrating data from a wide range of species enables insights into how millions of years of evolutionary processes have influenced cancer incidence, pinpointing molecular mechanisms of cancer suppression. Increasing the range of genomes used in comparative studies can reveal shared and lineage-specific variation across species, leading to the identification of novel mechanisms of cancer risks and resistance. For example, a cross-species reference genome analysis revealed how life-history traits such as lifespan and body size relate to cancer defenses [53]. Similarly, comparative genomic analyses across 63 mammalian species found that longer lifespan correlates with higher tumor suppressor gene copy numbers, indicating strong cancer defenses in long-lived species such as the naked mole rat [63]. Integrating cancer incidence data across many species, including whales, elephants and Tasmanian devils, can further reveal the molecular mechanisms underlying why certain species are notably vulnerable or resistant to cancer [10]. By leveraging data from many species, researchers can uncover evolutionary insights that may guide cancer treatment and prevention in humans.

There is currently no harmonized repository of mammalian cancer omics data spanning both human and non-human species. However, several notable human pan-cancer datasets have been instrumental in advancing cancer research. Large-scale cancer genomics initiatives such as The Cancer Genome Atlas (TCGA) [67], the International Cancer Genome Consortium (ICGC) [16], and the Pan-Cancer Analysis of Whole Genomes (PCAWG) [3] have generated extensive human cancer genomics datasets across multiple cancer types, including molecular data types such as gene expression, somatic mutations and copy number alterations. The Treehouse Childhood Cancer Initiative [8] is the largest compendium of childhood cancer data and highlights how transcriptomic data can be harmonized across many studies. Together, these resources demonstrate the value of large-scale harmonized cancer datasets.

One barrier to multi-species studies is the lack of cancer incidence and mortality data for most species [15, 66]. Recent efforts have focused on coordinating the sharing of biospecimens and associated data across zoological and conservation communities [34], as well as developing cross-species data portals such as the Global Genome Biodiversity Network (GGBN) Data Portal [19]. Additional efforts, such as the Arizona Cancer Evolution Center [13], highlight growing interest in cross-species cancer analysis. Cross-species transcriptomic databases such as Bgee [7] provide gene expression data from normal tissues.

Both this information and omics datasets spanning many species are essential to enable large-scale comparative cancer genomics across mammals. The lack of a centralized resource encompassing both human and non-human mammalian cancer data presents a significant challenge for multi-species comparative analysis.

Here we present the Paipu pipeline to facilitate the transition of comparative oncology studies from human plus one or two other species to pan-mammalian analyses. Paipu takes as input a list of species and search terms, then finds and retrieves all RNA-seq samples on SRA that fit those search terms. Paipu downloads, processes, and harmonizes metadata and gene expression data for all samples it retrieves. The resultant output is a ready-to-use atlas. We apply Paipu to 240 non-human mammals in the Zoonomia multispecies alignment [2] using 188 cancer-related keywords, creating a pan-mammalian tumor atlas that facilitates cross-species comparative transcriptomics.

## Materials and methods

The Paipu pipeline consists of three phases: (1) genome preparation, (2) SRA metadata retrieval and harmonization, and (3) RNA-seq data processing. The workflow is implemented in Nextflow [18] to manage processing across all three phases. In the genome preparation phase, Paipu downloads and preprocesses reference genomes for read mapping. During the SRA metadata retrieval phase, Paipu collects relevant cancer metadata and sequence data. Finally, in Paipu’s RNA-seq data processing phase, it uses the FREYA framework [26] to generate RNA-seq count data from the SRA raw read files.

Paipu’s output is organized by species. Each species directory includes a reference genome and annotation, harmonized metadata, BioProject information, and BioProject subfolders. The BioProject information file includes BioProject IDs, which are unique identifiers for each BioProject defined by SRA. It also includes sample IDs, sequencing layout, instrument platform, and submitter organization. BioProject subfolders are organized by sequencing layout (single-end or paired-end), each containing FASTQ files for all samples associated with that BioProject and layout type. Each layout subfolder also includes a configuration file, phenotype file, SRA accession file listing sample IDs, and RNA-seq count files. The Paipu pipeline steps are visualized in Figure 1 and each phase is described in detail below.

**Fig 1:**
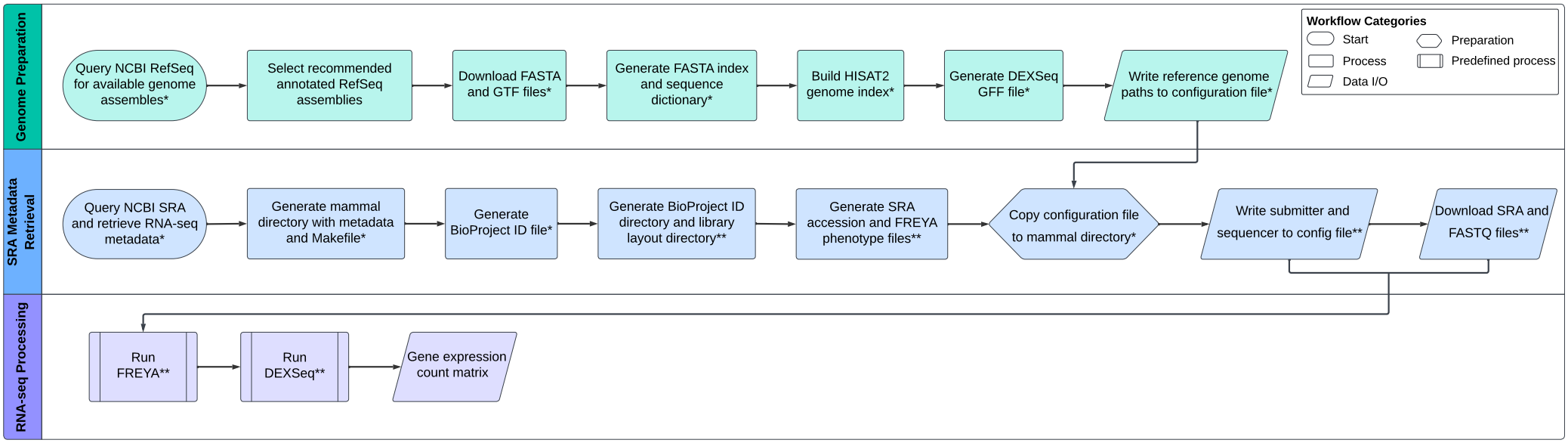
Overview of the Paipu framework. The workflow consists of three main components: reference genome preparation, involving downloading and preprocessing reference genomes for read mapping; SRA metadata retrieval, involving gathering essential metadata and sequence data; and RNA-seq processing via the FREYA pipeline. Asterisks indicate whether steps are performed per mammal (*) or per BioProject (**). Alt text: Workflow diagram of the Paipu framework showing three main components: reference genome preparation, SRA metadata retrieval, and RNA-seq processing via the FREYA pipeline, with asterisks denoting steps performed per mammal or per BioProject.

### Phase 1: Genome Preparation

For each species, Paipu performs a reference genome preparation phase that retrieves and organizes the required genome and annotation files. Paipu accepts an input file with two columns, one containing a list of species and the other containing a list of search terms. Paipu uses the species column to query available genome assemblies from NCBI using NCBI Datasets command-line interface [54] and identifies the current recommended reference genome based on assembly release date, annotation availability and associated metadata. It only prepares assemblies that have an associated gene annotation file.

For each assembly, Paipu downloads the genome FASTA and Gene Transfer Format (GTF) annotation files. Once the files are retrieved, the pipeline creates a genome sequence dictionary using Picard [1] and a corresponding FASTA index using samtools [41] to facilitate efficient random access to specific regions of the genome. The pipeline then builds a HISAT2 [36] genome index using the reference FASTA and GTF files to support RNA-seq read alignment. To detect differences in exon usage, it converts the GTF annotation into a General Feature Format (GFF) file using the DEXSeq [5] annotation preparation script. Paipu writes the paths of all generated reference and index files to a configuration file for use in the RNA-seq processing phase.

For species without an NCBI gene annotation, we incorporated TOGA [38] annotations after confirming that chromosome names matched those in the genome FASTA file. Users may incorporate alternative annotations following the same verification.

All reference genome assemblies were obtained from NCBI RefSeq [24] in September 2024 (see Table 1), with the exception of *Bos indicus* and *Tupaia chinensis*, which were obtained in November 2025. For *Enhydra lutris*, the reference genome assembly was retrieved from NCBI, while gene annotations were obtained from TOGA [38]. The reference genomes are organized into distinct mammal directories. Table 1 summarizes the genome assemblies and annotation sources used.

**Table 1.** Summary of reference genomes and sample counts per species in the pan-mammalian pan-cancer atlas.

| **Latin name** | **Common name** | **Clade** | **Ref genome version** | **Annot source** | **Protein-coding genes** | **Genome size (Gb)** | **Samples** |
|---|---|---|---|---|---|---|---|
| *Bos indicus* | zebu cattle | Cetartiodactyla | GCF_029378745.1 | NCBI | 20,976 | 2.7 | 16 |
| *Bos taurus* | domestic cattle | Cetartiodactyla | GCF_002263795.3 | NCBI | 21,667 | 2.8 | 38 |
| *Bubalis bubalis* | water buffalo | Cetartiodactyla | GCF_019923935.1 | NCBI | 21,359 | 2.6 | 3 |
| *Canis lupus familiaris* | dog | Carnivora | GCF_011100685.1 | NCBI | 21,175 | 2.5 | 2,262 |
| *Chlorocebus sabaeus* | green monkey | Primates | GCF_015252025.1 | NCBI | 21,620 | 2.9 | 1 |
| *Cricetulus griseus* | Chinese hamster | Rodentia | GCF_003668045.3 | NCBI | 21,776 | 2.4 | 28 |
| *Enhydra lutris* | sea otter | Carnivora | GCF_002288905.1 | TOGA | 19,126 | 2.5 | 1 |
| *Equus caballus* | horse | Perissodactyla | GCF_002863925.1 | NCBI | 20,995 | 2.5 | 16 |
| *Felis catus* | domestic cat | Carnivora | GCF_018350175.1 | NCBI | 20,453 | 2.4 | 11 |
| *Heterocephalus glaber* | naked mole-rat | Rodentia | GCF_000247695.1 | NCBI | 19,978 | 2.6 | 3 |
| *Macaca mulatta* | Rhesus monkey | Primates | GCF_003339765.1 | NCBI | 21,121 | 3.0 | 69 |
| *Mesocricetus auratus* | golden hamster | Rodentia | GCF_017639785.1 | NCBI | 21,616 | 2.5 | 28 |
| *Mus musculus* | house mouse | Rodentia | GCF_000001635.27 | NCBI | 22,198 | 2.7 | 48,077 |
| *Oryctolagus cuniculus* | rabbit | Lagomorpha | GCF_009806435.1 | NCBI | 22,458 | 2.8 | 12 |
| *Pan troglodytes* | chimpanzee | Primates | GCF_028858775.2 | NCBI | 22,561 | 3.2 | 1 |
| *Rattus norvegicus* | Norway rat | Rodentia | GCF_036323735.1 | NCBI | 23,209 | 2.8 | 938 |
| *Sus scrofa* | pig | Cetartiodactyla | GCF_000003025.6 | NCBI | 20,666 | 2.5 | 53 |
| *Tupaia chinensis* | Chinese tree shrew | Scandentia | GCF_000334495.1 | NCBI | 21,077 | 2.8 | 4 |
| **Total** |  |  |  |  |  |  | **51,561** |

### Phase 2a: SRA Metadata Retrieval

The SRA metadata retrieval phase of Paipu queries the NCBI SRA database [45] to identify RNA-seq samples that match the user-provided species and search terms, retrieves metadata for all matching samples using the NCBI Entrez Programming Utilities (E-utilities) [46] via Biopython [14] (version 1.79), which are subsequently harmonized. For each species in the input file, Paipu queries SRA for each search term provided to retrieve matching metadata. Each query combines the species and search term with built-in parameters specifying the sequencing strategy (`rna seq`), database name (`SRA`), and return type (`xml`). Metadata is retrieved in batches of 500, as recommended by the NCBI E-utilities [46], using Entrez esearch to identify matching records, Entrez read to extract unique identifiers (UIDs), and Entrez efetch to retrieve metadata. Paipu uses a short delay between requests to adhere to NCBI API usage guidelines. It parses retrieved records to extract metadata columns, then cleans and harmonizes the metadata. Paipu then combines data across search terms for each species and remove duplicate records based on run accession.

We applied this metadata retrieval approach as follows. To create the pan-mammalian pan-cancer atlas, we used a species list of 240 mammals from the Zoonomia consortium [2] and a manually curated a list of 188 search terms (Supplemental Table S1). These search terms cover a wide range of cancer types from the National Cancer Institute’s (NCI) list of cancer types [31] and the Surveillance, Epidemiology, and End Results (SEER) cancer statistics explorer network [56]. Together, these inputs enable retrieval of detailed metadata on experimental parameters, sequencing platforms, species, tissue types, and tumor classifications. Figure 2 shows a phylogenetic tree of all mammals from the Zoonomia alignment, annotated with tumor sample counts from the data atlas presented here. Figure 3 summarizes the tumor types and tissues represented in the pan-mammalian pan-cancer atlas.

**Fig 2:**
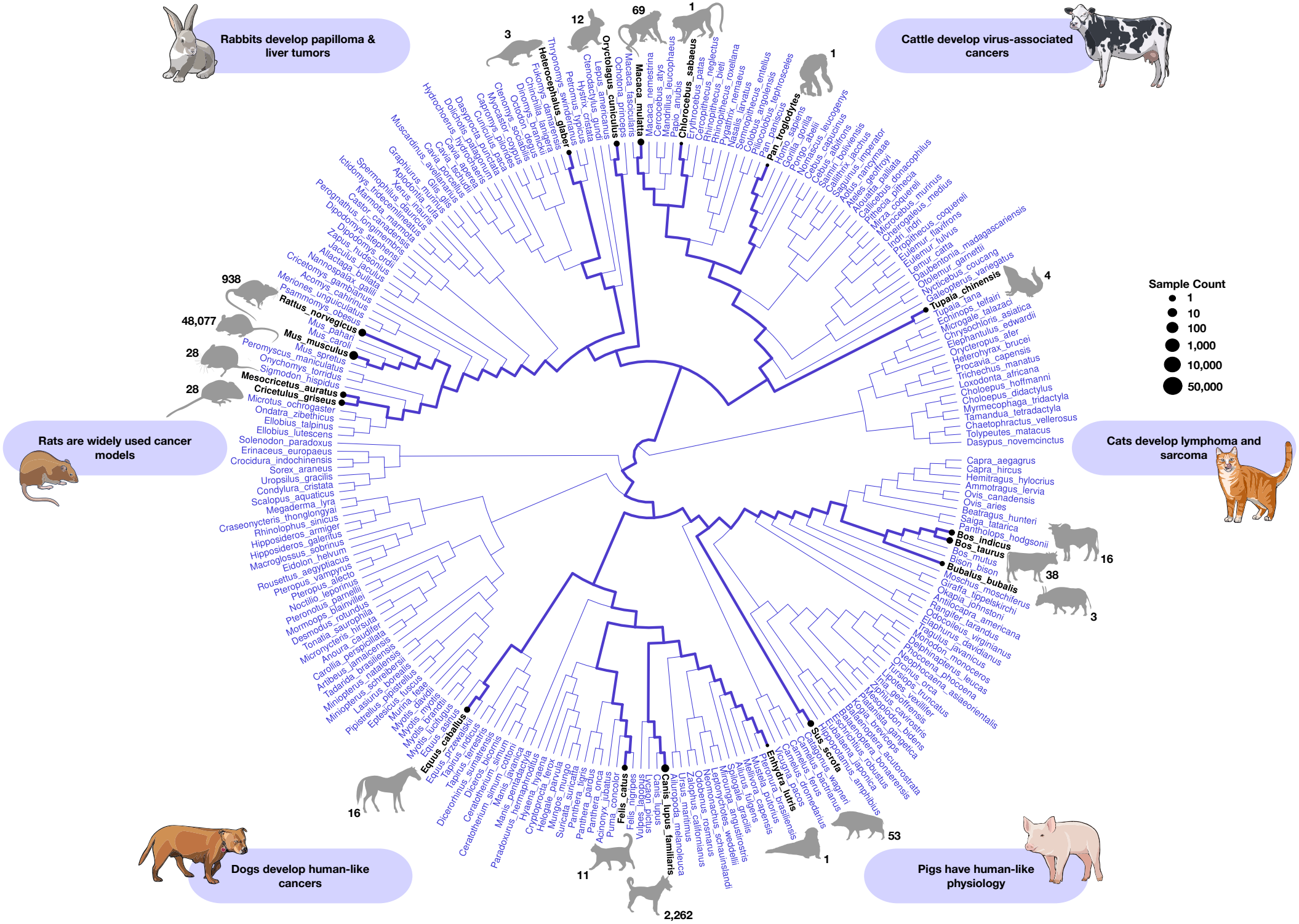
Phylogenetic tree of cancer samples across mammals. The phylogenetic tree of Zoonomia species shows the distribution of cancer samples across species. Black tip labels represent species with cancer samples and purple tip labels represent species without. Point size reflects log-scaled sample counts for species with data. Annotations highlight key cancer types or biological traits in selected species. Animal images were obtained from the NIAID NIH BioArt Source [47, 48, 49, 50, 51, 52]. Alt text: Phylogenetic tree of Zoonomia mammalian species showing the distribution of cancer samples across species. Species with cancer samples are labeled in black and species without samples are labeled in purple. Point sizes represent sample counts, and selected species are annotated with cancer types or biological traits.

**Fig 3:**
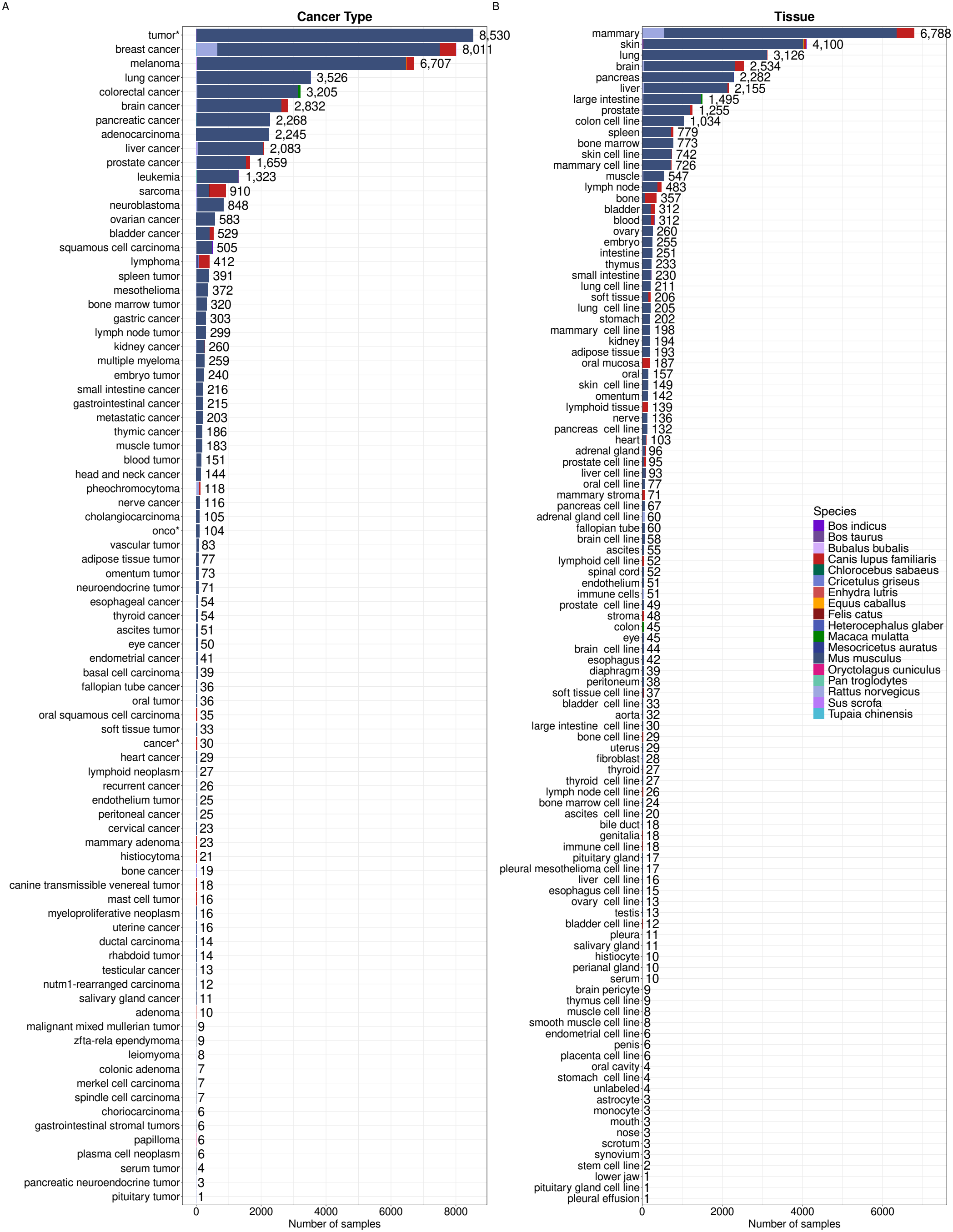
Biological metadata across species. Bar plots show biological and sample composition across species. (A) Cancer type and (B) tissue distributions are shown, where bars represent sample counts and colors represent species. Breast cancer is the most common cancer type and it is observed across *Mus musculus, Canis lupus familiaris, Rattus norvegicus, Tupaia chinensis* and *Bubalis bubalis*. A similar trend appears in the tissue distribution. Alt text: Bar plots showing biological metadata across species labeled A and B. A shows cancer type distributions and B shows tissue distributions, with bar heights representing sample counts and colors representing species. Breast cancer is the most common cancer type, with a similar pattern observed in the tissue distribution.

### Phase 2b: Metadata Harmonization

The metadata harmonization phase standardizes and cleans retrieved sample metadata to ensure consistency across studies. As previously noted [25, 39, 58], there is a non-negligible amount of spelling errors and other discrepancies that complicate harmonizing sample metadata. For example, we identified 5 spelling variants of the term ‘cell type’. To address spelling issues in the metadata, Paipu uses Levenshtein distance with a threshold of 3 to identify matching metadata column names with minor spelling differences, which are merged into one metadata column. This Levenshtein distance threshold was selected based on unit testing using a variety of thresholds on a sample set of metadata ranging from 2 to 8. We chose a threshold of 3 because it detected the most true positive errors. Overall, this approach improves metadata consistency by reducing redundancy.

We harmonized sequencing strategy by classifying samples as bulk or single-cell based on sample metadata. We labeled samples as single-cell if associated metadata columns contained any of the following keywords: ‘single-cell’, ‘single cell’, ‘scRNA-seq’, ‘scRNAseq’, ‘scRNA’ and ‘singlecell’. We classified all remaining samples as bulk.

After metadata retrieval and initial harmonization by Paipu, we performed additional post-processing steps to standardize metadata across studies further. We standardized common missing metadata value labels (e.g. ‘nan’, ‘unknown’, ‘missing’) to NA across all metadata columns and removed repeated words and duplicated phrases within metadata values. We harmonized several metadata attributes (e.g. cell type, tissue, tumor grade, treatment, breed) by combining information from related metadata columns. Keyword and pattern matching on metadata column names identified related columns for each metadata attribute to be combined. The first non-empty value across these related columns was used to create a single harmonized metadata column.

We standardized sex labels by combining information from multiple sex-related metadata columns and mapping them to MALE, FEMALE, or UNKNOWN. We also standardized sequencing platform annotations and labeled missing values as ‘Unknown’. We standardized tissue annotations further by using regular expression (regex) patterns for tissue names and related terms, including synonyms, abbreviations, spelling variations and cancer-related terms. When a tissue could not be identified using the regex patterns, we inferred it from cancer type annotations or other metadata columns, such as source names, when possible. Samples without an identified or inferred tissue were labeled as ‘unlabeled’.

We further standardized cancer type annotations using tissue information and other metadata columns, including sample source name, cell type, disease, and tumor type. We used keyword and pattern matching within these columns to infer missing cancer type annotations when possible. For samples with generic cancer annotations assigned based on the search term used to identify the sample (e.g., ‘cancer*’, ‘tumor*’, ‘‘onco*’), we inferred cancer type from the harmonized tissue annotation when available. We manually assigned tissue annotations to 2,704 samples that remained unlabeled after pattern matching and corrected cancer type annotations for 52 BioProjects. We then grouped cancer types into broader cancer systems using a predefined mapping derived from NCI Cancers by Body Location/System classification [30]. As a final step, metadata columns that consisted entirely of missing values were removed.

These harmonization steps reduced the total number of metadata columns from 263 to 212 in the non-mouse dataset and from 1,334 to 841 in the mouse dataset. All post-processing steps were implemented in R outside of Paipu to allow for specific atlas harmonization without modifying the framework. We caution that incomplete or ambiguous metadata and terms not identified by keyword or pattern matching may lead to errors during harmonization.

#### Quality control and filtering

To produce a high-quality atlas, we implemented a methodical process of quality control and filtering. Our original Paipu search found 671,352 relevant samples. Subsequent analysis of these data identified a critical issue in the SRA search function. Samples are returned when a search term appears in any metadata column, including the organization name. As a result, the initial search results included many samples that were unrelated to cancer but were processed at Cancer Centers. To address this, we added a post-search filtering step to the Paipu pipeline. After downloading metadata but before downloading raw reads or processing the samples, Paipu performs a secondary filtering process and removes samples that do not meet the user-provided inclusion criteria. In this post-filtering step, Paipu removes samples in which search terms appear only in metadata columns describing submitter, organization or sequencing center information. Samples are kept if the search term also appears in any other metadata column.

#### Data organization

After downloading and harmonizing the sample metadata, Paipu organizes this information into an easily digestible structure. It then downloads the RNA sequencing reads associated with each SRA accession number in the metadata, in preparation for data processing. To bin samples, Paipu uses the BioProject information file to create a directory for each BioProject. This directory contains subdirectories for ‘single’ and ‘paired’ layout reads. Each subdirectory stores the corresponding SRA accession numbers in a file. Using the SRA accession file, a phenotype file is created in CSV format which contains each SRA accession, a sequentially assigned identification number and a ‘T’ for tumor sample classification for FREYA [26], given that only tumor samples are included.

For the configuration setup, Paipu copies the configuration file associated with each species to its BioProject layout subdirectories and populates each copy with BioProject metadata, including submitter and sequencing platform information. The configuration files are used by FREYA [26] in the RNA-seq processing phase. Paipu then retrieves the raw sequencing data using tools such as prefetch and fastq-dump from the NCBI SRA Toolkit [23] to download the SRA files and convert them into FASTQ format. This structured setup enables efficient, organized access to RNA-seq data for each species.

### Phase 3: RNA-seq Processing

The RNA-seq processing phase uses FREYA [26], a framework for RNA-seq data processing and analysis. FREYA performs read alignment using HISAT2 [36], quality control with FastQC [6], and quantifies gene-level expression with DEXSeq-Count [5, 59]. It removes duplicate reads with MarkDuplicates [1], splits and adjusts spliced reads with SplitNCigarReads [64] and assigns reads to alleles with AddOrReplaceReadGroups [1]. For each BioProject layout, the Paipu pipeline schedules FREYA to perform read alignment, quality control and expression profile generation. For each sample, FREYA generates high-quality counts files. Once FREYA is complete for a given mammal, Paipu runs DEXSeq [5] to aggregate gene-level counts for each BioProject layout and subsequently generates a count matrix for each. Paipu then combines these counts data into a single gene expression matrix file for each BioProject layout.

### Optional second-pass filtering using a metadata classifier

To classify samples as cancer or non-cancer, we developed a metadata-based classifier that integrates regex pattern matching with clinical natural language processing (NLP). Free-text metadata columns, including sample source name, tissue, disease and study title, are used for classification. The classifier uses a multi-stage approach to identify terms associated with cancer and non-cancer.

First, sample metadata is scanned using regex patterns for more than 300 terms associated with cancer and non-cancer, including ‘melanoma’, ‘carcinoma’, ‘normal’ and ‘healthy’. Cancer and non-cancer signals are then identified and used to assign an initial regex classification. Sample specific information take priority over broader study information during classification. Next, sample metadata is evaluated using MedSpaCy [22] to incorporate contextual information and detect negation. This allows phrases such as ‘no evidence of tumor’ and ‘non-cancerous’ to be classified as non-cancer. MedSpaCy evaluates each selected metadata column separately. Columns that contain terms associated with cancer are recorded as supporting evidence for the assigned classification label.

The regex and MedSpaCy classifications are combined to assign a final classification and confidence label. When classifications conflict between the two methods, sample-level evidence is prioritized over study-level evidence. For example, a sample identified as normal tissue is classified as non-cancer even when the study title contains terms associated with cancer. Based on the combined results, samples are classified into one of five categories: ‘confident_cancer’, ‘likely_cancer’, ‘confirmed_by_medspacy’, ‘confirmed_non_cancer’ or ‘likely_non_cancer’. The classification categories are then mapped to a final cancer or non-cancer label based on their respective classification. These categories are also grouped into ‘confident’ or ‘likely’ for downstream analyses.

After cancer classification, sample metadata is scanned using regex patterns to identify benign and cell line samples. More than 50 cell line patterns are included, consisting of known cell line names (e.g.,’B16’, ‘CT26’) and terms such as ‘cell line’. Benign patterns include terms such as ‘benign’ and ‘non-malignant’.

The final cancer or non-cancer classifications and confidence levels were added to the harmonized metadata as ‘paipu_cancer_label’ and ‘paipu_cancer_confidence’, respectively. Benign and cell line flags were added as ‘paipu_is_benign’ and ‘paipu_is_cell_line’, respectively. Samples were not removed based on these classifications and flags because public metadata may be incomplete or ambiguous.

### Statistical analysis of metadata

We performed statistical tests in R to assess associations between selected metadata variables. Associations between categorical metadata variables were evaluated using Pearson’s Chi-square tests. For contingency tables with small expected counts, we estimated p-values using Fisher’s exact test with Monte Carlo simulation (100,000 replicates). We compared quantitative metadata columns across species using the Kruskal-Wallis test. Sequencing read counts were estimated from NCBI SRA total spot counts based on library layout, with 2 reads per spot for paired-end sequencing and 1 read per spot for single-end sequencing. Forty-one *Mus musculus* samples with missing total spot counts were excluded from the sequencing read analysis. To evaluate the relationship between malignancy prevalence and sample count, we calculated sample counts from our atlas metadata and merged them with malignancy prevalence estimates from Compton et al [15], excluding species without available estimates (n=3). Multiplicity adjustment was performed using the Benjamini-Hochberg false discovery rate procedure.

### Sex prediction using transcriptomic data

We predicted sex for samples with missing sex metadata using gene expression from the full joined non-mouse expression dataset and the *Mus musculus* expression dataset separately. Gene expression data was normalized using the trimmed mean of M-values (TMM) method in edgeR [60] and transformed to log2 counts per million (logCPM) with a prior count of 1. We defined sex-specific markers using *XIST* as a female marker and a set of Y chromosome genes as male markers, including *DUSP9, CD99, THOC2, SMARCA1, TMSB4Y, RPS4Y1, DDX3Y, EIF1AY, KDM5D, UTY, USP9Y* and *ZFY* [32]. Y chromosome expression was calculated as the sum of expression across these genes for each sample.

We evaluated the predictive performance of *XIST* and Y chromosome gene expression individually using receiver operating characteristic (ROC) analysis on samples with known sex labels and calculated the area under the curve (AUC). Females were defined as the positive class. We then trained a logistic regression model with female sex as the outcome and *XIST* expression and aggregated Y chromosome expression as predictors to evaluate their combined performance.

The combined model had the strongest predictive performance. We applied this model to all samples to obtain the probability of being female and assigned predicted sex labels using a threshold of 0.5. For samples with known sex labels, we kept the original values, but for samples without sex labels, we assigned the predicted labels. We defined prediction confidence based on predicted probabilities, with high confidence for probabilities *≥* 0.9 or *≤* 0.1, medium confidence for probabilities between 0.75–0.9 or 0.1–0.25 and low confidence otherwise.

### Principal component analysis

We performed principal component analysis (PCA) to explore patterns in cancer gene expression across mammals (Figure 4A,B). For all PCA analyses, we restricted the expression data to only include genes shared across samples. For non-mouse samples, samples with *<*2,600 detected genes were removed (see Supplemental Figure 1A). For *Mus musculus*, samples with *<*2,000 detected genes were removed (see Supplemental Figure 1B). The non-mouse and *Mus musculus* datasets were processed separately. Genes with low expression were filtered using edgeR [60], with groups defined as the interaction between species and cancer type. After filtering, the gene expression data was normalized using the TMM method in edgeR and transformed to logCPM with a prior count of 1. For non-mouse samples, we calculated pairwise Pearson correlations among samples within each species to reduce redundancy, and samples with correlation *>*0.99 were filtered by removing the sample with the higher mean correlation to other samples (see Supplemental Figure 2). The deduplicated non-mouse dataset was combined with the *Mus musculus* dataset to form the final atlas.

**Fig 4:**
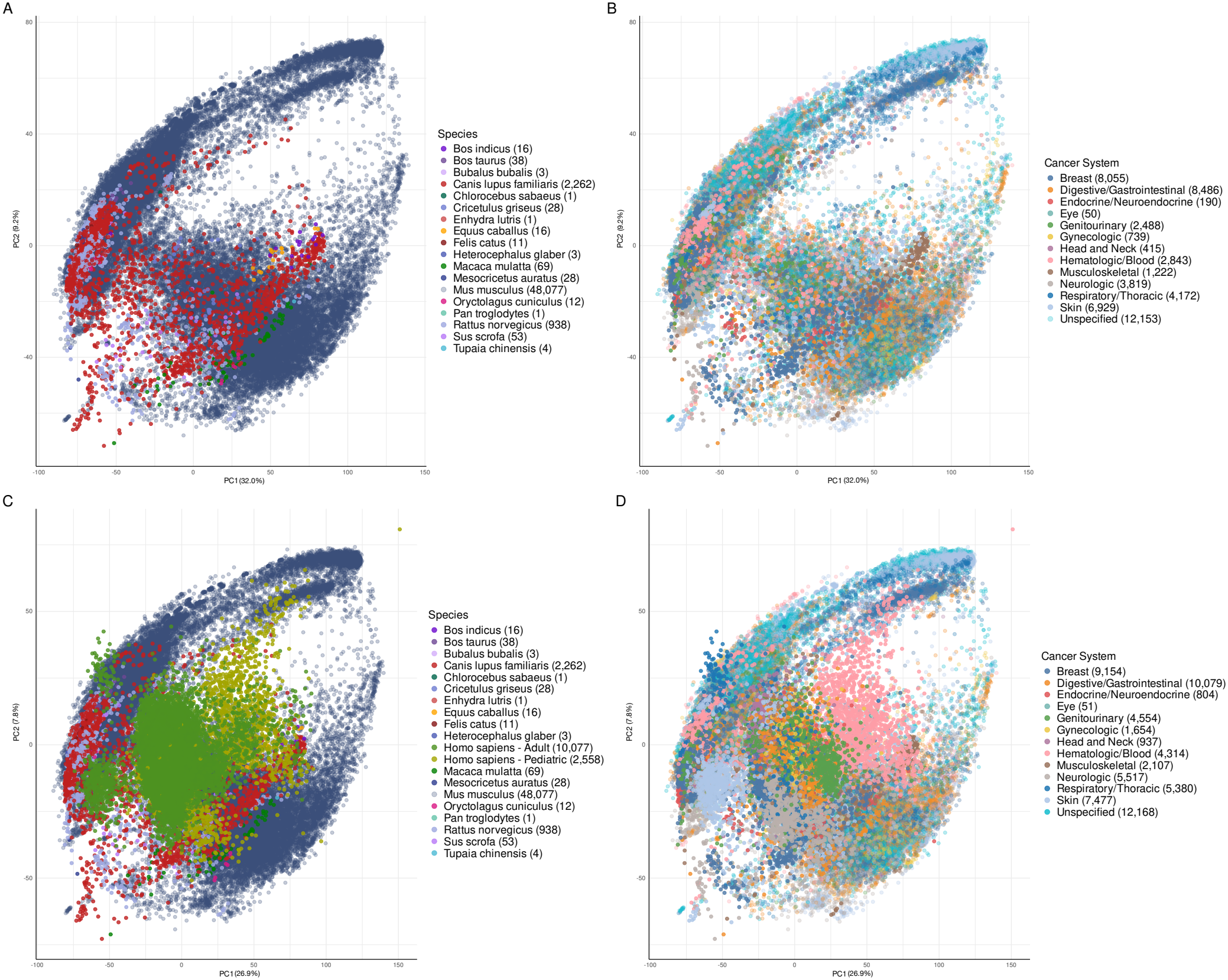
Principal component analysis (PCA) of cancer gene expression across the pan-mammalian pan-cancer atlas with and without the Treehouse dataset. (A and B) PCA of the atlas, with samples colored by species (A) and cancer system (B). (C and D) PCA of the atlas and Treehouse dataset, with samples colored by species (C) and cancer system (D). Sample counts are shown in parentheses in each legend. Alt text: Principal component analysis plots of gene expression across the pan-mammalian pan-cancer atlas labeled A and B. Principal component analysis plots of gene expression across the pan-mammalian pan-cancer atlas and the Treehouse dataset labeled C and D. A and C are colored by species and B and D are colored by cancer system. Sample counts are shown in parentheses in each legend.

**Fig 5:**
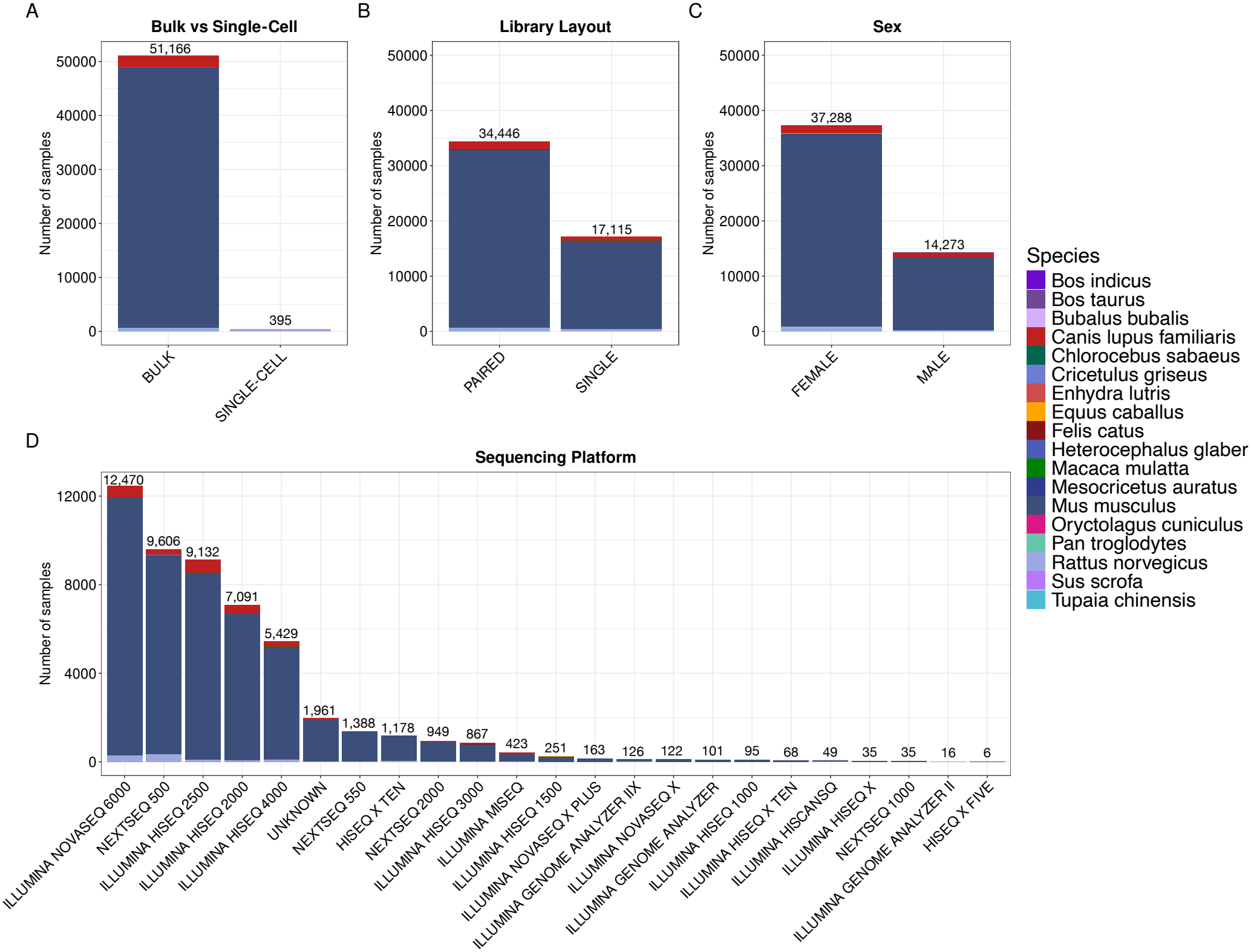
Technical metadata across species. Bar plots show the distribution of key technical and sample attributes across species. (A) Bulk versus single-cell sequencing, (B) library layout, (C) sex, and (D) sequencing platform are shown, where bars represent sample counts and colors represent species. Bulk sequencing and paired-end libraries are most common. Illumina platforms are used for most samples. Alt text: Bar plots showing technical metadata across species labeled A to D. A shows bulk versus single-cell sequencing, B shows library layout, C shows sex and D shows sequencing platform distributions. Bar heights represent sample counts and colors represent species. Bulk sequencing, paired-end libraries, female samples and Illumina platforms are most common.

We conducted PCA on our final atlas and visualized results with samples colored by species and by cancer system. We then extended the analysis to include human samples from the Treehouse compendium [8]. We excluded samples with unknown ages and classified human samples as pediatric (age *≤* 18) or adult (age *>* 18). Genes with zero expression across all samples were removed before normalization. The dataset was then normalized using TMM method in edgeR [60] and transformed to logCPM with a prior count of 1 to be consistent with the processing applied to the final atlas. The normalized Treehouse and atlas datasets were combined by shared genes and batch-corrected between human and non-human samples using ComBat [33]. We then performed PCA on the combined dataset and visualized the results with samples colored by species and by cancer system (Figure 4).

### Evaluating species robustness as models of human cancers

As a case study for the use of the Paipu data atlas, we extracted all species-cancer pairs with at least 7 samples from the pan-mammalian pan-cancer atlas and applied the species-cancer robustness formula from Daluwatumulle et al [17] to quantify how well each species models each human cancer, where data is available. Supplemental Table S2 shows sample counts for all combinations that met our inclusion criteria. To assess whether a species is a robust model organism for a given cancer, we applied the formula and adjusted the cutoff from Daluwatumulle et al [17] to this new dataset (Figure 6). We statistically tested whether certain mammalian species tend to score higher more often than expected by chance, using a permutation test. Scores were used to rank species-cancer combinations and the mean rank was then calculated for each species. The analysis was repeated using 10,000 permutations, in which species labels were randomly shuffled and mean ranks recalculated. For each species, we calculated a one-sided p-value by comparing the observed and the shuffled mean ranks. Multiplicity adjustment was performed on all p-values using the Benjamini-Hochberg false discovery rate procedure. This analysis was performed separately for adult and pediatric comparisons. We also calculated statistical differences between human and non-human species gene expression, using the Mann-Whitney U test. For this analysis, we selected the top 10 most frequently mutated genes from the COSMIC database.

**Fig 6:**
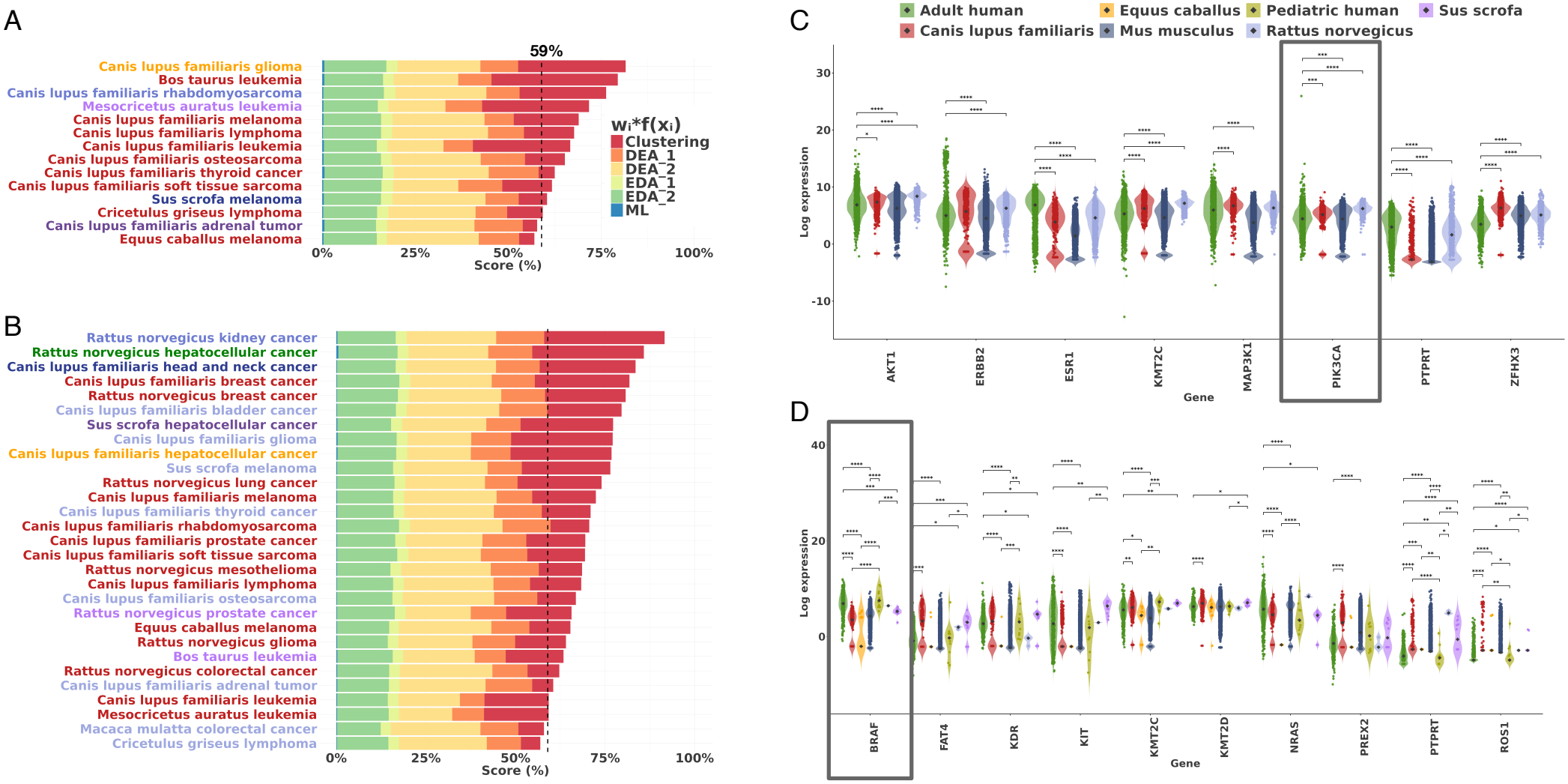
Identifying robust transcriptomic models of human cancers using the Paipu data atlas. (A-B) Species-cancer model scores using the formula from Daluwatumulle et al [17] to rank species according to their transcriptomic similarity to (A) pediatric and (B) adult cancers. Color coding indicates the contributions of the formula components for each score. The vertical dashed lines show the robustness cutoff value for these data. Scores greater than this value are considered robust cancer-species combinations. (C-D) Violin plots with jittered points showing log-scaled gene expression values in COSMIC genes associated with (C) breast cancer and (D) melanoma for species with at least 7 samples, colored by species. Median expression values are marked as a diamond. Differences in gene expression between human and non-human species were assessed for each cancer type using the Mann-Whitney U test. Statistical significance is shown as follows: \**p ≤* 0.05, \*\**p ≤* 0.01, \*\*\**p ≤* 0.001, \*\*\*\**p ≤* 0.0001. Alt text: Bar charts of formula results ranking species according to their similarity to pediatric and adult cancers labeled A and B, respectively. Gene expression distributions of breast cancer and melanoma are labeled C and D, respectively.

## Results

### Pan-mammalian pan-cancer atlas

We used Paipu to create a pan-mammalian pan-cancer atlas from our manually curated list of cancer terms and mammalian species from the Zoonomia multi-species alignment [2]. Human data was excluded from this publication. Paipu identified and retrieved data for 4,753 non-mouse mammalian cancer RNA-seq samples spanning 20 species, along with 51,288 mouse (*Mus musculus*) bulk cancer RNA-seq samples. Using the 188 search terms increased samples significantly. For example, Cahill et al [12] identified 1,437 dog tumor samples using the search term ‘cancer’, whereas Paipu identified 2,835 dog tumor samples. Filtering to remove samples in which cancer-related keywords were only present in non-biological metadata columns reduced this atlas to 4,112 samples spanning 19 species. After duplicate removal, this corresponded to 3,484 samples spanning 17 species (Figure 2). *Mus musculus* samples were processed separately, with 48,077 samples remaining after filtering low-quality samples. A final atlas, combining mouse and other species data, contained 51,561 samples. Samples with available run size information (n=51,534) represented 98.5 TB of sequencing data, with individual run sizes ranging from 0.5 MB to 94.0 GB (median 1.4 GB, mean 2.0 GB). Figure 3 shows the cancer types and tissues found for each species. Rodentia was the most represented clade in the data atlas, followed by Cetartiodactyla, Carnivora and Primates. As of April 30, 2026, 219 species had no cancer-related RNA-seq data in SRA.

### Atlas composition

The atlas contains multiple tumor types across species and various sequencing characteristics. Some tumor types were more frequent in certain species; for example, osteosarcoma samples were predominantly from *Canis lupus familiaris* (n=327), while 54 samples were from *Mus musculus* and 1 was from *Rattus norvegicus*. Breast cancer was the most common tumor type, followed by melanoma, lung cancer and colorectal cancer (Figure 3A). Lymphoma and leukemia were the most widely represented tumor types, occurring in 7 and 6 species, respectively.

Samples with available estimated read counts (n=51,520) had a median of 43.1 million reads per sample ranging from 0.02 *−* 1, 976.4 million reads. Sequencing type also differed by species. Single-cell RNA-seq was represented in *Rattus norvegicus* (38.5%), *Heterocephalus glaber* (100%) and *Canis lupus familiaris* (1.4%), whereas the remaining non-mouse species were represented by bulk RNA-seq. Only bulk RNA-seq samples were included for *Mus musculus*. Sequencing type was significantly associated with library layout (Pearson’s Chi-squared test with Yates’ continuity correction, adjusted *p* = 1.06 *×* 10^*−*28^).

Metadata harmonization improved annotation consistency across samples. For non-mouse samples, the number of unique cancer type labels was reduced from 55 to 35 by aggregating related cancer subtypes into broader categories (e.g. sarcoma subtypes), while retaining the original subtype annotations in a separate column. Tissue annotations were combined into a single column through automated harmonization and manual curation. This tissue harmonization step increased the proportion of samples with tissue annotations from 67% to 100%. Additionally, the proportion of samples with missing biological sex annotations decreased from 59% to 0% following prediction from gene expression. For *Mus musculus* samples, the number of unique cancer type labels was reduced from 122 to 70 by aggregating related cancer subtypes into broader categories. Tissue harmonization increased the proportion of samples with tissue annotations from 63% to 67%, while prediction from gene expression decreased the proportion of mouse samples with missing biological sex annotations from 80% to 0%.

### Metadata-based sample classification

The metadata-based classifier was evaluated using 410 manually annotated non-mouse samples. Overall accuracy was 90.7%, with precision 98.3% and recall 89.8% for cancer samples. Among the 3,484 non-mouse samples, 2,434 (69.9%) were classified as cancer and 1,050 (30.1%) as non-cancer. In total, 3,050 (87.5%) classifications were ‘confident’, while 434 (12.5%) were ‘likely’. 2,905 (83.4%) samples were flagged as cell lines and 190 (5.5%) as benign. Among the 48,077 mouse samples, 38,986 (81.1%) were classified as cancer and 9,091 (18.9%) as non-cancer. In total, 36,066 (75.0%) classifications were ‘confident’, while 12,011 (25.0%) were ‘likely’. Additionally, 21,260 (44.2%) samples were flagged as cell lines and 49 (0.1%) as benign.

### Principal component analysis and integration

After filtering samples based on the number of detected genes, the non-mouse dataset was reduced from 4,112 to 3,935 samples. Removing highly similar samples based on sample correlation further reduced the non-mouse dataset to 3,484 samples. For *Mus musculus*, filtering samples based on the number of detected genes reduced the dataset from 51,288 to 48,077 samples.

To evaluate variation in the expression data, we performed PCA on the combined non-mouse and *Mus musculus* RNA-seq data (Figure 4A,B). The samples followed a continuous distribution in PCA. When colored by species, there was species-specific grouping, which is consistent with differences in gene expression across mammals. When colored by cancer system, there was partial grouping of related cancer systems. These results indicate that species is a major driver of variation and that cancer-related signal is also present.

To evaluate integration with external datasets, we integrated the atlas with the Treehouse compendium [8], including TCGA data, and performed PCA on the combined dataset (Figure 4C,D). The samples followed a continuous distribution in PCA, with a different structure compared to the Paipu atlas alone. Additionally, samples from the pan-mammalian pan-cancer atlas and Treehouse compendium overlapped within cancer systems. These results indicate that the processed expression data is comparable across datasets.

### Individual project enrichment

This pan-mammalian cancer atlas contains data from 3,507 BioProjects and 51,561 samples. BioProjects have on average 14.70 samples (median 9.0, IQR 6.0-15.0, SD 27.45), demonstrating the high variability in study sizes. The combined data atlas is 3,508 times larger than the average BioProject.

### Statistical enrichment in species metadata

The pan-mammalian tumor atlas consists primarily of *Mus musculus* and *Canis lupus familiaris* samples, consistent with expectations given their use in medical and basic science settings. Together, these two species make up 97.6% of the atlas, with 93.2% *Mus musculus* and 4.4% *Canis lupus familiaris* samples (Figure 2). Across all species, most samples are bulk RNA-seq (see Figure 5A, 99%). Most samples are sequenced using paired-end reads (67%), while 33% use single-end reads (Figure 5B). More than 20 unique sequencing platforms are represented in the atlas, with Illumina NovaSeq, HiSeq, and NextSeq used most frequently (Figure 5D).

We then assessed the relationship between cancer type and species using harmonized cancer type categories that aggregate related cancer subtypes. Cancer type was significantly associated with species (Fisher’s exact test with Monte Carlo simulation, adjusted *p* = 1.25 *×* 10^*−*5^). Library layout was also significantly associated with species (Fisher’s exact test with Monte Carlo simulation, adjusted *p* = 1.25 *×* 10^*−*5^), and sequencing reads showed significant differences across species (Kruskal-Wallis test, *χ*^2^ = 272.77, *df* = 17, *p* = 2.27 *×* 10^*−*47^). In contrast, we found no evidence of a monotonic association between malignancy prevalence and sample count across species (Spearman’s *ρ* = 0.16, *p* = 0.57).

### Predicting sex from transcriptomic data

Given that information about biological sex was only available for 41% of non-mouse duplicated samples, we sought to determine the sex of unlabeled samples using computational techniques. Based on results from previous studies [32], we evaluated the predictive performance of *XIST* and aggregated expression of Y chromosome genes for sex classification using samples with known sex labels (1,052 female and 547 male). ROC analysis showed that aggregated expression of Y chromosome genes alone achieved an AUC of 0.60, while *XIST* expression alone achieved an AUC of 0.85. A logistic regression model with female sex as the outcome and *XIST* expression and aggregated expression of Y chromosome genes as predictors achieved an AUC of 0.93. Using a probability threshold of 0.5, the combined logistic regression model achieved an accuracy of 88.7%, with sensitivity of 92.2% and specificity of 81.9%. We then applied the logistic regression model to the 2,336 unlabeled non-mouse tumor samples. The model predicted 1,584 samples as female and 752 samples as male.

For *Mus musculus*, the same approach was evaluated using samples with known sex labels (6,643 female and 3,199 male). Aggregated expression of Y chromosome genes alone achieved an AUC of 0.66, while *XIST* expression alone achieved an AUC of 0.75. The combined logistic regression model achieved an accuracy of 76.6%, with sensitivity of 88.2% and specificity of 52.2%. We then applied the model to the 38,295 unlabeled samples, predicting 28,294 samples as female and 10,001 samples as male. After incorporating predicted labels for both non-mouse and *Mus musculus* samples, the pan-mammalian pan-cancer atlas contains no missing biological sex labels and consists of 72% female and 28% male samples (Figure 5C).

### Identifying robust mammalian models of human cancers

As a case study, we applied the formula from Daluwatumulle et al. [17] to evaluate the robustness of each species as a transcriptomic model for human cancer (Figure 6A,B). Excluding mouse, there was sufficient data to analyze 59 species-cancer combinations, including 19 adult human and 11 pediatric human cancers. Of these, 37 species-cancer combinations scored above the robustness cutoff (59%). The permutation test identified pigs and dogs as statistically significant candidate models for adult cancers (both adjusted *p* = 1.28 *×* 10^*−*2^), and domestic cattle as a statistically significant candidate model for pediatric cancers (adjusted *p* = 1.20 *×* 10^*−*3^).

We selected breast cancer and melanoma (Figure 6C,D) to explore the expression distribution across species. We selected these two cancers because they met three criteria: they ranked among the top 10 unique cancer types exceeding our cutoff, included data for more than two non-human species, and both have approved drugs, as reported by Mateo et al. [44]. The expression profile of drugs targeting the *BRAF* gene show no statistically significant differences between human (adult and pediatric) and rat samples (Figure 6D), while *PIK3CA* shows inter-species variability across all species (Figure 6C). Both genes also show high variability in widely used animal models such as *Mus musculus*.

### Pipeline Performance

We ran Paipu on the University of Florida’s HiPerGator high-performance computing cluster using the SLURM scheduler. Processing reference genomes and RNA-seq data and associated metadata across 20 species (4,753 samples) required approximately 1,600 hours of wall time and approximately 38,000 core hours of compute. We processed the *Mus musculus* initial dataset (51,288 samples) using similar resource allocations. The compute estimate is based on allocated cores and runtime, rather than actual CPU usage.

## Discussion

Despite the increasing volume of available RNA-seq data, cross-species cancer studies are still limited by the lack of unified datasets that span many species. Here, we developed Paipu, a pipeline that finds and harmonizes SRA data from many species relating to a user-provided list of species and keywords. We apply Paipu to create a pan-mammalian pan-cancer transcriptomic atlas. This resource enables cross-species analyses that were previously difficult due to inconsistent metadata and data being spread across studies.

The pan-mammalian tumor atlas presented here highlights the scale of publicly available transcriptomic data and the challenges associated with integrating many datasets from different sources. Paipu initially identified 4,753 non-mouse samples across 20 species. After filtering, quality control, and duplicate removal, 3,484 samples spanning 17 species remained. This reduction in size highlights the importance of quality control when integrating public data. An additional 51,288 *Mus musculus* bulk RNA-seq samples were identified, with 48,077 samples remaining after filtering and quality control. Altogether, the final atlas contains 51,561 samples spanning 18 species. It contains over 98.5 TB of sequencing data across 3,507 studies. The atlas also reveals strong biases in data availability, as most of the samples are from a small number of species, particularly *Mus musculus, Canis lupus familiaris*, and *Rattus norvegicus*. For most of the 241 mammals in the Zoonomia alignment, there is no cancer RNA-seq data in NCBI SRA.

Some tumor types are more frequent in certain species, reflecting both biological differences and sampling biases. For example, we found that osteosarcoma samples were predominantly from *Canis lupus familiaris*, which is consistent with its higher incidence rate in dogs compared to humans [12]. In addition to biological differences, sequencing methods also contribute to this data bias. Single-cell RNA-seq data differs from bulk RNA-seq in the amount of information it captures and is often more sparse due to dropout. Dropout occurs when a gene is present in a cell, but is not detected because only small amounts of its mRNA are captured during sequencing [57, 70]. The limited representation of single-cell RNA-seq data across species is likely due to the increased technical complexity of single-cell RNA-seq, higher costs [69], challenges in sample preparation [62] and sensitivity to incomplete annotations, especially in non-model organisms [28]. Only 395 single-cell samples exist in the pan-mammalian data atlas, most of which are *Rattus norvegicus* (rat) tumors. Of these, most (n=363) are breast cancer samples.

Technical differences were observed in sequencing platforms, library layouts and the number of reads per sample. These differences can introduce biases, though approaches such as batch correction may help reduce some of these effects. Some variability is expected, however, given diversity in experimental designs and sequencing strategies across the 3,507 studies. This highlights the importance of approaches that enable consistent downstream processing of large integrated datasets.

A major challenge in using publicly available data is incomplete and inconsistent metadata annotation [39, 58], which may limit the accuracy of metadata harmonization and downstream analyses. Paipu includes a filtering layer and a metadata-based sample classifier that can be used as a second filtering layer. However, some annotation errors may remain despite automated harmonization and manual curation. In some cases, analysis of sequencing data can be used to fill metadata gaps. For example, biological sex information was spread across 12 metadata columns. Paipu harmonization created a global biological sex column using these data, in which 77% of samples were still missing sex labels (n=40,631). To address this, we implemented a logistic regression model based on *XIST* and aggregated Y chromosome gene expression. This model successfully identified biological sex for all samples, resulting in zero missing data for the sample sex column and thus a more complete atlas. This demonstrates how missing information can be deduced from genomic data using computational techniques to address common challenges related to incomplete metadata in public datasets.

Paipu facilitates creation of high-quality datasets, enabling comparative transcriptomics across many species. As a case study, we evaluated gene expression in two cancers with known drug-gene interactions involving drugs used in their clinical treatment. According to the similarity formula, dog tumors ranked closer to human tumors than rat tumors in breast cancer, whereas, pigs, dogs and horses ranked closest to adult human tumors, in that order, in adult melanoma. We found that the expression distributions of *PIK3CA*, the target of the approved drug alpelisib for breast cancer, and *BRAF*, the target of the approved combination therapy of atezolizumab, vemurafenib and cobimetinib for BRAF V600E/K-mutant melanoma, showed varying degrees of overlap between human tumors and non-human mammalian species. Understanding these similarities and differences is important for strengthening the translation of preclinical drug research into clinical applications.

While existing tools encompass similar individual Paipu components, they do not provide a comprehensive resource. MetaSRA [9] provides normalized metadata for human samples from the NCBI SRA, addressing challenges in metadata standardization. MetaSRA is, however, limited to human samples. Harmonized data atlases such as Recount3 [68] include more studies than presented here, but these data are restricted to human and mouse samples. The NextFlow nf-core/rnaseq workflow [21] is an alternative for the gene expression data processing and could replace the FREYA pipeline currently used by Paipu. While flexible and well-designed, this workflow does not process metadata.

The absence of a centralized resource that integrates reference genome preparation, metadata retrieval and RNA-seq processing for non-human mammals remains a significant gap in the field. Paipu addresses this gap by enabling consistent processing of RNA-seq data and associated metadata across studies in NCBI SRA. By systematically retrieving and harmonizing public datasets from different studies, Paipu facilitates the use of data that would otherwise remain separate and difficult to compare. It improves the usability of public datasets and facilitates cross-species analyses.

## Acknowledgments

Support was provided by NIH NCI R01 #R01CA265907 and #R01CA265907-03S1. This project makes use of publicly available data from the Sequence Read Archive. Graphical abstract icons were adapted from resources provided by Streamline (https://streamlinehq.com).

## Author contributions statement

Bria S. Smith (Data curation [], Formal analysis [], Investigation [], Methodology [], Software [], Validation [], Visualization [], Writing–original draft [equal], Writing–review & editing [equal]), Leslie A. Smith (Data curation [], Investigation [], Software [], Validation [], Visualization [], Writing–review & editing [equal]), Geesa Daluwatumulle (Formal analysis [], Investigation []), Nathan Glen (Software [], Methodology [], Validation []), Ji-Hyun Lee (Formal analysis [], Validation [], Writing–review & editing [equal]), James A. Cahill (Conceptualization [], Methodology [], Supervision [], Formal analysis [], Funding acquisition [], Validation [], Writing–review & editing [equal]), Kiley Graim (Conceptualization [], Formal analysis [], Funding acquisition [], Investigation [], Methodology [], Software [], Supervision [], Validation [], Visualization [], Writing–original draft [equal], Writing–review & editing [equal])

## Supplementary Data

Supplementary data is available at *NAR* online.

## Data availability

All sequencing data and associated metadata used in this study were obtained from publicly available data from the NCBI Sequence Read Archive. We provide seven dataset versions, including raw unfiltered, processed and deduplicated datasets, along with their harmonized metadata. The processed datasets, including the pan-mammalian pan-cancer atlas are available on Zenodo under https://doi.org/10.5281/zenodo.24066813. The raw unfiltered datasets are available under https://doi.org/10.5281/zenodo.24067794. Code for Paipu, along with scripts for data processing and downstream analyses, can be accessed at https://github.com/GraimLab/Paipu, and code for the metadata-based sample classifier can be accessed at https://github.com/NathanG1en/Paipu-Metadata-Classifier.

## Supplemental Information

**Table S1.**

240 organisms and 188 cancer-related search terms used as input to Paipu to query and retrieve RNA-seq samples.

**Table S2.**

Sample counts across 19 cancer types and 10 mammalian species included in the case study.

**Supplemental Figure 1:**
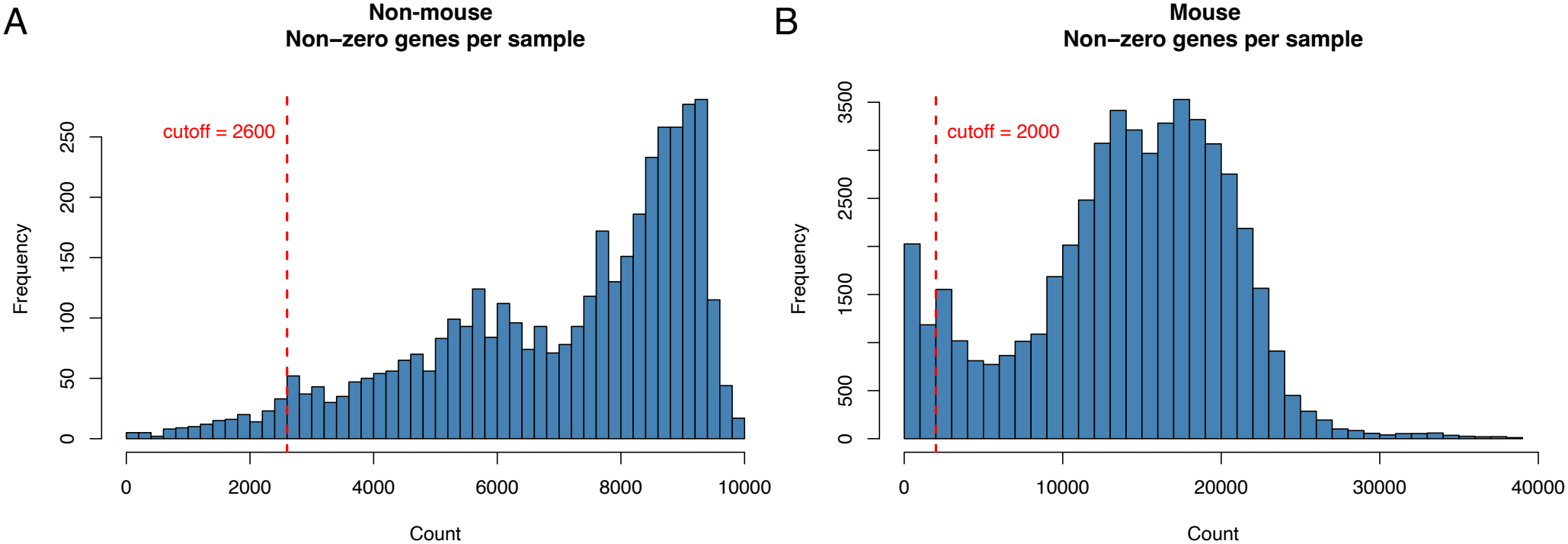
Distribution of expressed genes per sample. Histogram of the number of expressed genes (non-zero expression) per sample. (A) Non-mouse samples. Samples with *<*2,600 expressed genes were excluded as low-quality. (B) Mouse samples. Samples with *<*2,000 expressed genes were excluded as low-quality. Cutoffs were chosen based on the respective distributions of expressed genes per sample. Alt text: Histograms of the number of expressed genes per sample, labeled A and B. A shows the distribution for non-mouse samples, with a cutoff at 2,600 expressed genes. B shows the distribution for mouse samples, with a cutoff at 2,000 expressed genes.

**Supplemental Figure 2:**
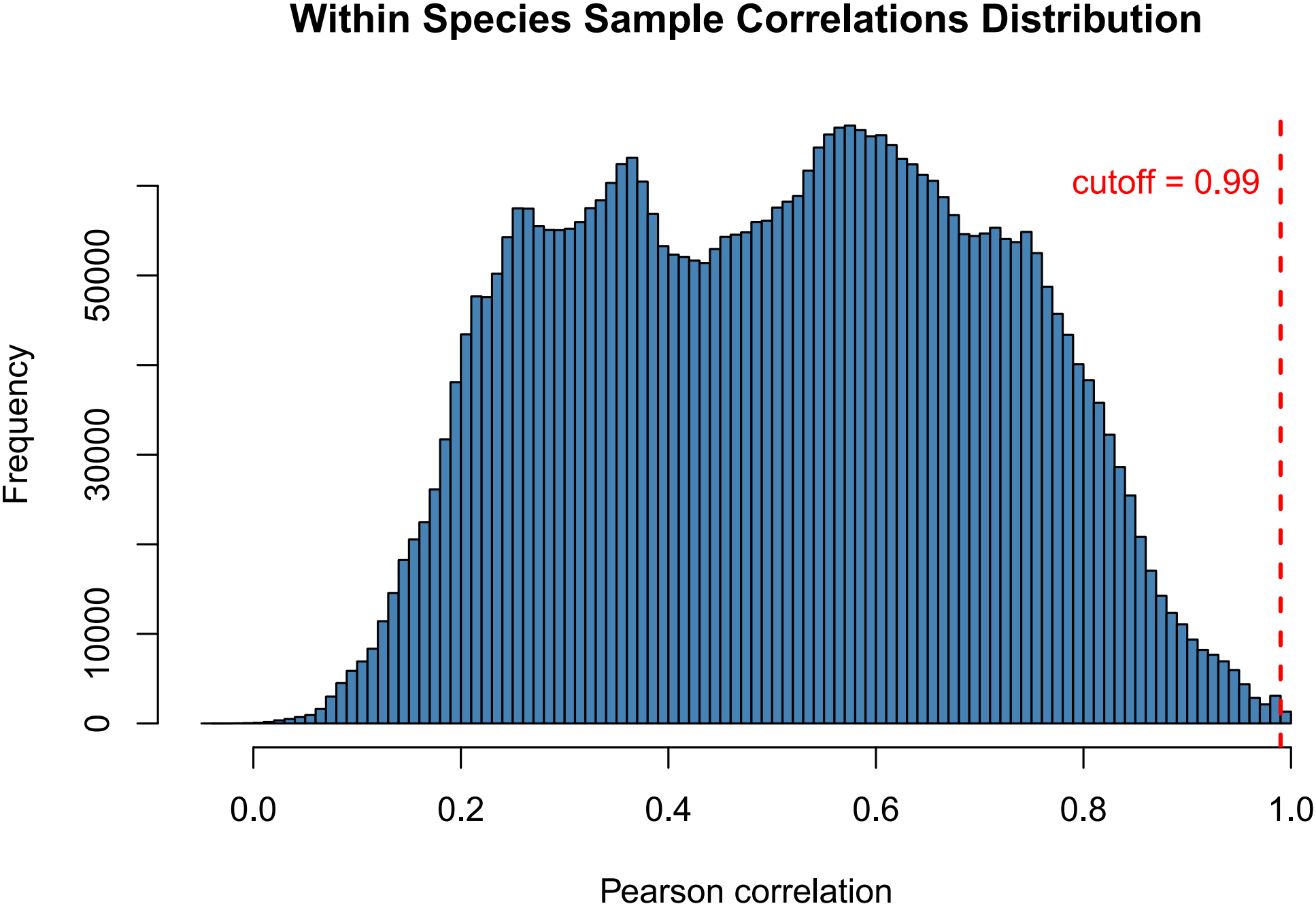
Distribution of within species sample correlations. Histogram of within species pairwise Pearson correlations. Sample pairs with correlation *>*0.99 were considered highly correlated and one sample from each pair was removed based on its mean correlation to other samples. Alt text: Histogram of within species pairwise Pearson correlations between samples. A cutoff at 0.99 indicates highly correlated sample pairs.

